# A Highly Efficient Aorta-Gonad-Mesonephros-Like Definitive Hemogenic Endothelium From Human Pluripotent Stem Cells

**DOI:** 10.1101/2023.06.26.546545

**Authors:** Yesai Fstkchyan, Qingwen Cheng, Jingli Zhang, Daniel Lu, Guanyi Huang, Tiange Dong, Lauren Jones, Matt Kanke, Chris Hale, Kristin Tarbell, Chi-Ming Li, Songli Wang, Stuart M. Chambers

**Affiliations:** Research Biomics, Amgen Research, South San Francisco, CA; Genome Analysis Unit, Amgen Research, South San Francisco, CA; Oncology Research, Amgen Research, South San Francisco, CA

## Abstract

Human pluripotent stem cells are a tremendous tool to model early human development and disease including their use in the *in vitro* generation of blood cell fates. Hematopoietic progenitors and stem cells are the primary source of blood and the immune system from early development to adulthood and arise through successive waves of hemogenic mesoderm either in the yolk sac or embryo proper. Researchers have long sought a tractable human model for observing and distinguishing these waves of hematopoiesis in the dish for human developmental and disease modeling. Here we report a high-efficiency method for differentiating human pluripotent stem cells into an aorta-gonad-mesonephros-like definitive hemogenic mesoderm capable of giving rise to definitive hematopoietic progenitor and stem cells. The hematopoietic progenitor and stem cells exhibit robust multilineage *in vitro* colony forming potential. Gene expression analysis and single cell sequencing strongly support the developmental timing and notion that the pluripotent stem cell derived hematopoietic stem and progenitors are strikingly like *bone fide* hematopoietic stem cells. The hematopoietic progenitors can be subsequently differentiated into polarized macrophage and T-cells *in vitro*. Minimal silencing was observed upon differentiation of the pluripotent stem cells to hematopoietic lineages when conducting gene editing. Finally, upon engraftment into immunodeficient animals the hematopoietic progenitors and stem cells differentiate into multiple lineages including B-cells, T-cells, NK-cells, and monocytes.

## Introduction

Pluripotent stem cells hold tremendous promise in modeling human development and disease. They can be derived from either the inner cell mass of a human blastocyst, called human embryonic stem cells (hESCs), or via reprogramming of human somatic cells into pluripotent tissue, called induced pluripotent stem cells or iPSCs^1^. Since the initial derivation of pluripotent stem cells^2^, it is speculated both hESCs and iPSCs can give rise to any cell type in the human body, permitting a genetically tractable, limitless source of human tissue for drug development and cell therapy.

Hematopoietic progenitors and stem cells (HPSCs) create the blood and immune system for life and sit at a nexus for human disease and disease progression. Atop this hierarchy, hematopoietic stem cells (HSCs) self-renew and are responsible for the lifelong multipotent production of all blood cell fates^3^, including lymphocytes, myeloid cells, red blood cells, and platelets. HSCs are critical for the normal function of the immune system and the maintenance of hematopoietic homeostasis.

While the lineage tracing and developmental origin of blood are greatly complicated by the nature of a liquid tissue, evidence from mouse and iPSC studies suggests there are at least 4 independent waves of hemogenic mesoderm (HM) contributing to the blood as the embryo develops^4^: primitive (yolk-sac) hematopoiesis, erythromyeloid progenitor (EMP), lympho-myeloid primed progenitor cell (LMPP), and the long-term hematopoietic stem cell (LT-HSC). The aorta-gonad-mesonephros (AGM) is a mesodermal developmental region where LT-HSCs, capable of long-term multilineage engraftment (LT-HSCs) in adults first emerge. Human iPSCs permit scientists a window into early human development and are well suited for distinguishing between various waves of blood formation and there has been a tremendous effort to differentiate or reprogram LT-HSCs from either hESCs or iPSCs^5–12^ with most seminal reports indicating great strides towards making an aorta-gonad-mesonephros (AGM) -like HPSC production^11, 13^, with the hallmark of increased lymphocyte production^14^. However, more work is needed in teasing apart cell fate decisions since studies have failed to achieve long-term, multilineage engraftment.

During human development, LT-HSCs emerge from a hemogenic endothelium by undergoing an endothelial-to-hematopoietic transition (EHT), marked by the expression of Runt-related transcription factor 1 (RUNX1), CD34, and CD45, and the downregulation of the endothelial markers SRY-Box Transcription Factor 17 (SOX17) and CD31. Knock-out studies of the RUNX1 locus in rodents, also called AML1, exhibit normal yolk sac hematopoiesis and embryonic lethality at E12.5 from a lack of fetal liver hematopoiesis^15^. Importantly, studies that focus specifically on the C spliceoform suggest it plays a role in LT-HSC specification, self-renewal, and differentiation^16^, and is required for proper hematopoietic differentiation from iPSCs^17^. More efficient protocols allowing observation of cell fate decisions in a discrete and stepwise fashion are necessary to move the field forwards.

Our group has strived to improve efficiencies of cell fate derivation through lessons learned from prior work on the high-efficiency derivation of the nervous system^18^ applied to hematopoietic lineages with the goal of making progress towards generating LT-HSCs without using cellular aggregation (embryoid bodies or organoids) or gene delivery, often used for cell fate reprogramming^19^. Here we report a high-efficiency method for generating a HM with all the molecular and many functional hallmarks of a definitive AGM-like wave of hematopoiesis including synchronous EHT, robust expression of TBXT and RUNX1C, single-cell gene expression signature clustering with *in vivo* counterparts for a population of iPSC derived cells, and robust generation of myeloid and lymphoid cell fates *in vitro*. The method is amendable to gene editing, even though gene silencing is prevalent within pluripotent stem cell differentiation and is capable of multilineage hematopoiesis persistent for up to 12 weeks *in vivo*.

## Results

### HSPCs undergo EHT and emerge between days 14 and 21 in vitro

The directed differentiation was divided into four discrete stages based on the natural development of the AGM (**Figure 1A, B**): mesoderm specification, endothelium specification, EHT, and HPSC expansion and differentiation. By day 9 of cultures, endothelium formation was observed based on morphology and the expression of CD31 (not shown). Between days 14 and 21, phase bright free-floating cells began to emerge indicating the transition of EHT and hematopoiesis (**Figure 1B**). The harvested suspension cells when examined by flow cytometry were found by sequential gating to be 77% Lin-, 49% CD34/CD45 positive, 8% CD38/CD45RA negative, and 80% CD90 positive. A heterozygous iPSC reporter line engineered to express eGFP from the distal RUNX1C promoter was engineered to quantify the emergence of a definitive wave of hematopoiesis using the same method. The emergence of eGFP positive cells were observed in blood islands between days 14 and 21, corresponding with EHT. Upon further examination by flow cytometry, within the CD34/CD45 positive fraction, nearly all cells were eGFP positive indicating a robust production of definitive wave of HPSCs.

**Figure 1:**
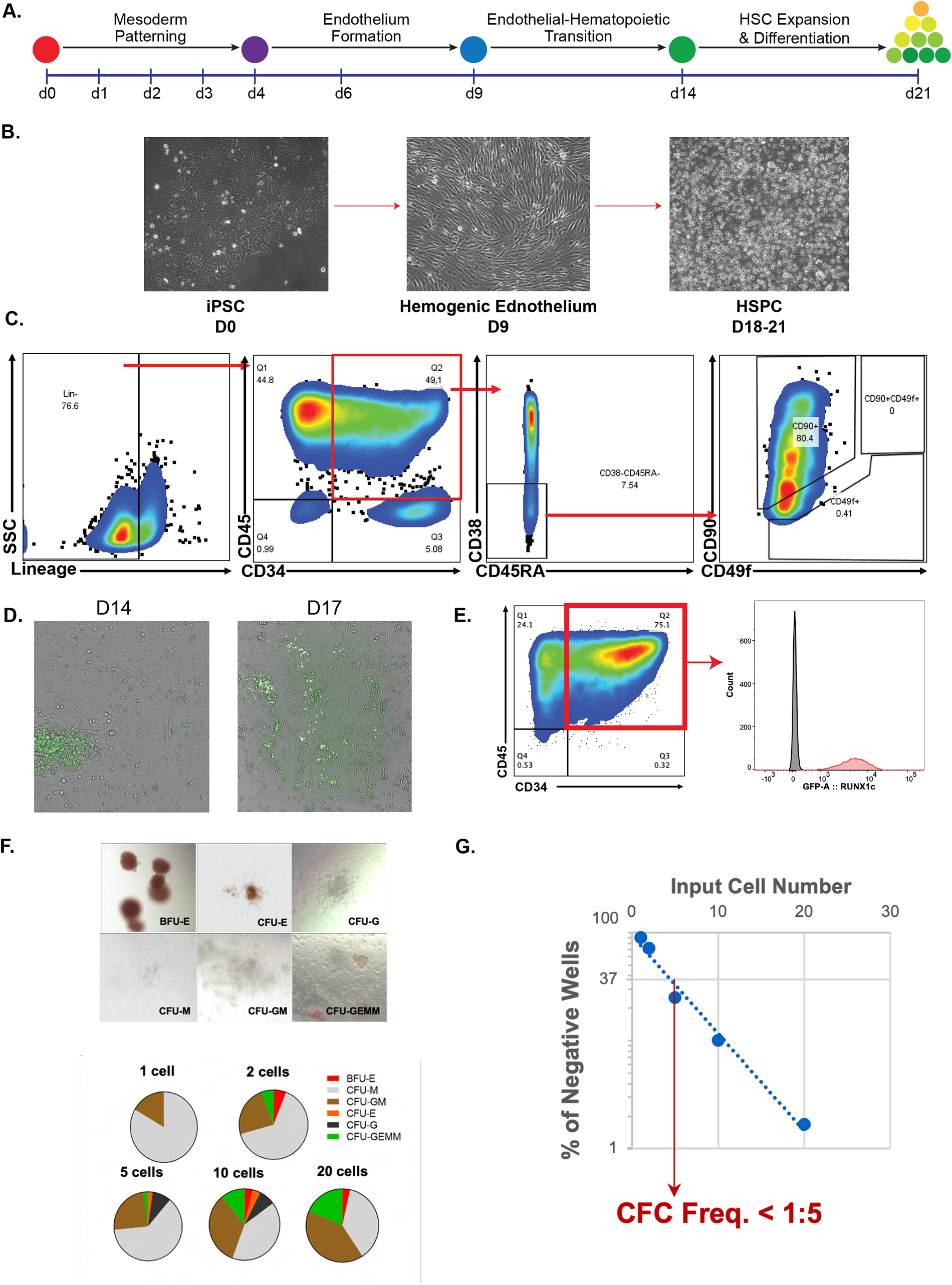
A. Schematic of the differentiation of LT iPSC to definitive hematopoietic stem cells (HSCs). In short, iPSCs were plated as single cells and then inducted to undergo developmental-stage specific differentiation by first mesoderm induction, followed by endothelial formation. Lastly, a specified cocktail was added so that differentiation cultures undergo endothelial-to-hematopoietic transition and subsequent maturation. B. Brightfield images of day 0 (DO) iPSCs, hemogenic endothelium on day 9 (D9), and floating iPSC derived HSCs on day 18-20 (D18-20). C. Representative flow cytometry plots show the gating strategy for identifying HSCs from iPSC differentiation cultures. Here, we characterize HSCs as lin-CD45+CD34+CD45RA-CD38-/lowCD90+ D. Brightfield images overlayed with GFP channel showing green fluorescence in budding cells undergoing EHT. E. Representative flow cytometry plot showing the RUNX1c GFP expression in CD34+CD45+ cells from differentiation cultures F. Schematic describing the methylcellulose and limiting dilution assays to assess the potency of the iPSC derived HSCs. G. Representative brightfield images of various colonies found in CFU assays. Colonies were counted and categorized as BFU-E, CFU-E, CFU-G, CFU-M, CFU-GM, and CFU-GEMM. Quantifications are shown as both pie charts and bar graphs. H. Quantification of CFUs at various input cell numbers for limiting dilution analysis. CFC frequency was calculated based by calculating the number of cells necessary to achieve colonies in 37% of wells or ∼1 in 5 cells.

A limiting dilution methylcellulose colony forming unit (CFU) assay was performed to serve as the gold standard *in vitro* functional test of HPSCs and from 1 to 20 individual CD34/45/90 positive, CD38/Lin negative cells were sorted into a 96-well plate. CFU-granulocyte, erythrocyte, monocyte, and megakaryocyte (CFU-GEMM) colonies could be observed in wells containing as few as two sorted cells (**Figure 1F**) and an overall colony forming frequency was found to be one in five cells having CFU potential within this sorted fraction (**Figure 1G**) indicating a high prevalence of HPSCs in the cultures.

### EHT occurs via a TBXT-positive mesodermal intermediate

A qRT-PCR time course was performed to define the timing and stages of development in the dish to better understand the transition from mesoderm to endothelium. Markers for each stage were observed in sequential time windows corresponding to the morphological changes observed: mesoderm (days 2-4), endothelium and arterial (days 4-9), and AGM markers (days 4-14). According to a recent study, six reprogramming factors are sufficient to induce an HSC-like state^9^, perhaps indicating their role in supporting the HSC state by a transcription factor network. Of the six factors, we detected expression of ERG, HOXA5/9/10, and SPI1, but no LCOR expression was observed.

Moreover, the T-box transcription factor T (TBXT) is required for normal mesoderm and blood development in rodents^20^ and its expression appears at gastrulation, and is expressed in and required for posterior and axial mesoderm^21^. Additionally, recent reports have shown that extra-versus intra-embryonic mesodermal lineages from iPSC differentiations can be further discriminated the expression the cell surface markers CXCR4 and KDR/FLK1^22^, whereas single positive CXCR4 cells give rise to extra-embryonic mesodermal lineages and those cells expressing both CXCR4 and KDR/FLK1 are indicative of intra-embryonic mesoderm.

We next sought to further assess the developmental origin of our iPSC derived HSCs by examining the expression for the cell surface markers KDR/FLK1 and CXCR4, which have been shown to distinguish between intra- and extra-embryonic mesodermal lineages^22^. As definitive hematopoiesis arises from an intra-embryonic mesodermal lineage, we examined whether the mesodermal cells at day 3 of differentiation express both these markers (**Figure 2B**). We find that approximately 40% of cells have co-expression for CRXR4 and KDR/FLK, suggesting that most cells have the potential to give rise to definitive mesodermal lineages.

**Figure 2:**
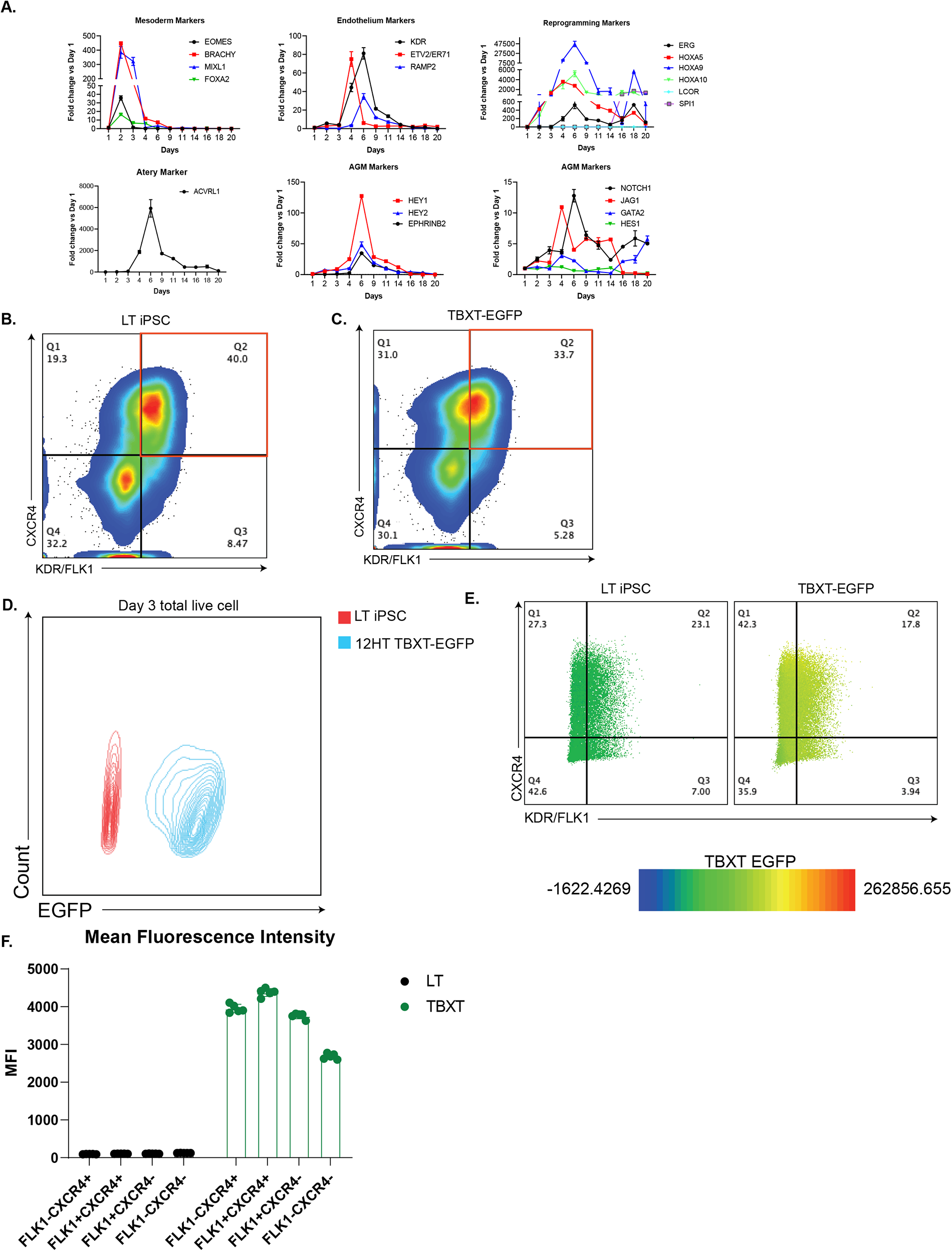
A. qRT-PCR analysis of mesoderm-, endothelial-, artery-, reprogramming-, and AGM-specific genes through the course of iPSC differentiation towards a definitive HSC. B. Representative flow cytometry plot of day 3 (D3) iPSC differentiation cultures measuring the expression of cell surface markers CXCR4 and KDR/FLK1. C. Day 3 expression of CXCR4 and KDR/FLK1 from the differentiation of the TBXT EGFP reporter line D. Expression of TBXT EGFP (red) at day 3 of differentiation compared to the parental LT iPSC line (blue) E. Representative flow cytometry plots at day 3 of differentiation utilizing both the LT and TBXT EGFP line showing expression on cell surface markers CXCR4 and KDR/FLK1. Shading of the plots displays EGFP expression intensity within the four specified populations. F. Quantification of the median fluorescent intensity within CXCR4-KDR/FLK1-, CXCR4+KDR/FLK1-, CXCR4+KDR/FLK1+, and CXCR4-KDR/FLK1-populations.

To further confirm our observations, we sought to generate a reporter iPSC line for TBXT to assess whether the expression of CXCR4 and KDR/FLK1 do indeed specify intra-embryonic mesoderm. To do so, we used genetic engineering to introduce a H2B-EGFP construct at the most distal polyadenylation signal site (PAS), immediately following exon 8 of the endogenous TBXT locus (**Figure S1C**). We then utilized this iPSC line to measure the expression of this lineage priming transcription factor within the CXCR4 and KDR/FLK1 populations. First, we ensured that the engineered line was able to faithfully differentiate to mesoderm by collecting cells on day 3 post differentiation. We then measured the emergence of intra-embryonic mesoderm by flow cytometry. We found that approximately 33.7% of cells were double positive for CXCR4 and KDR/FLK1 (**Figure 2C**). These results were comparable to the parental from which this line was generated from. Moreover, almost all the cells during this time point of differentiation express EGFP, suggesting TBXT transcriptional activity was present in all four mesodermal populations (**Figure 2D**). This is further confirmed by overlaying the EGFP intensity onto the four different mesodermal populations that are present on day 3 of differentiation (**Figure 2E**). Quantification of the TBXT-EGFP mean fluorescent intensity (MFI) revealed that the FLK1-CXCR4-population contained the least amount of TBXT reporter activity, whereas the CXCR4+FLK1+ population was amongst the highest (**Figure 2F**). Taken together, these data suggest that the use of CXCR4 and FLK1/KDR does not necessarily discriminate between intra- and extra-embryonic mesoderm as we observe TBXT reporter activity in four populations. Moreover, these data provide further evidence that our differentiation protocol faithfully directs iPSC differentiation cultures to an intra-embryonic mesodermal lineage, thus producing definitive iPSC derived HSCs.

### Single-cell sequencing reveals a population of iPSC derived HSPCs with striking similarities to bone fide hematopoietic stem cells

To further validate that our iPSC derived HSCs reflect the transcriptional program of those found in humans, we utilized single-cell RNA (scRNA) sequencing to compare the transcriptomes of the various stages of differentiation and the trajectory towards that of *bone fide* HSC from fetal liver (FL), umbilical cord blood (UCB), and bone marrow (BM). To do so, we sorted cells during specific differentiation time points based on adherence and cell surface markers (**Figure 3A**). Endothelial cells were isolated as adherent cell types based on the marker CD31, whereas AGM-like cells were identified as CD34+CD45-. Moreover, Pre-HSC populations were characterized as still adherent, but expressing the cell surface markers CD34 and CD45. Lastly, HSCs were identified by CD34+CD45+CD90+ and the more differentiated multipotent progenitors (MPP) by the lack of CD90 expression (CD34+CD45+CD90-). To further aid in the proper calculation of developmental trajectories, CD34-CD45+ cells were included in the experiment to represent those cells that have undergone additional differentiation steps toward terminal cell types of the blood.

**Figure 3:**
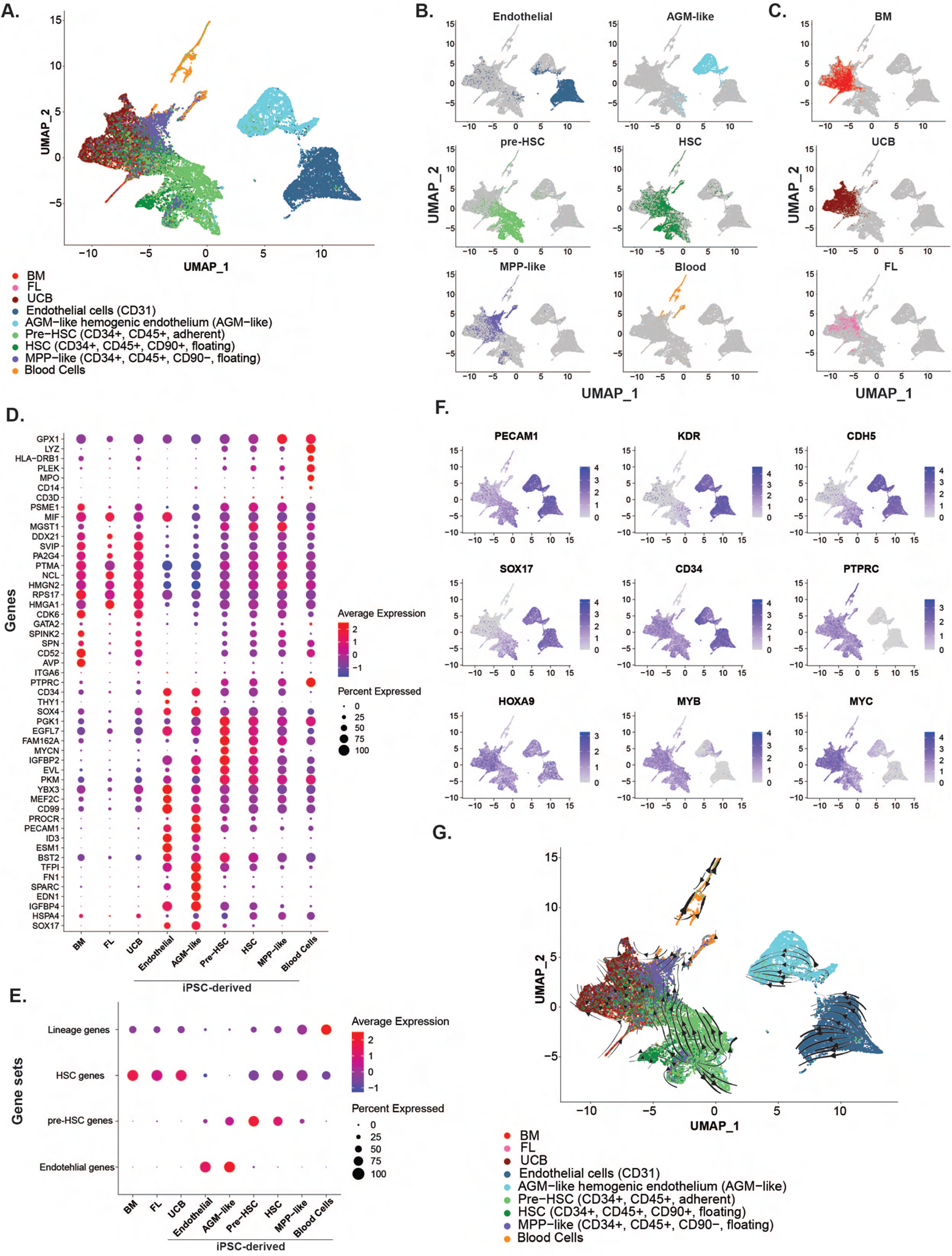
A. UMAP projection of the diffusion map (dmap) dimensionality reduction embeddings of the top 45 dimensions of the various induced pluripotent stem cell (iPSC) derived endothelial, hemogenic endothelium aorta-gonad-mesonephros (AGM)-like, pre-hematopoietic stem cell (pre-HSC), HSC, multipotent progenitor (MPP)-like, and blood cells. Additionally, human fetal liver (FL), umbilical cord blood (UCB), and bone marrow (BM) derived *bona fide* HSC’s were also included in he analysis. These data represent approximately 25% of that data (n=32,754 cells). F-statistic variance modeling was used to filter out noisy or lowly expressed genes, yielding 11,804 features that are stably expressed. Cells were sorted based on morphology, cell attachment, and hematopoietic markers (CD34, CD45, and CD90). B. Single population highlights of UMAP projection for the various iPSC derived populations throughout the course of differentiation. C. *Bona fide* human HSCs from three independent tissue sources. Each population contains biological and technical triplicates. D. Dot plot displaying the expression of key stage-specific hematopoietic and developmental genes across specified iPSC derived clusters and *bona fide* HSCs. Color represents the average expression, and the size of the dot indicates the percent of cells within the cluster expressing specified genes. E. Dot plot of cell-type identification through gene expression utilizing Seurat’s module score feature. Expression of gene modules is scored based on the average expression and percent of the population that expresses the set of given genes. F. Feature plots displaying the expression of key hemogenic genes established by the field that is expressed during distinct stages of development. G. UMAP projection of the RNA velocity was calculated using spliced and unspliced counts to solve transcriptional dynamics.

After data integration and clustering, we find that these isolated cell types represent distinct stages of differentiation (**Figure 3B**). Both endothelial and AGM-like cells cluster independently from Pre-HSCs and HSC populations, suggesting that a developmental transition has occurred in between. This is most likely characterized by the fact that hemogenic endothelial cells undergo an endothelial-to-hematopoietic transition (EHT) from an adherent cell type to suspension (ref). Furthermore, iPSC derived pre-HSCs and HSCs cluster closely together, suggesting a shared transcriptional profile, whereas the MPP population and blood cells cluster away from HSCs. Interestingly, *bone fide* HSCs from BM, UCB, and FL all cluster together and in approximately the same dimensions as our iPSC derived HSCs, suggesting that our iPSC derived HSCs are transcriptionally related to *bone fide* HSCs (**Figure 3C**).

To ascertain the shared transcriptional profiles of iPSC derived HSCs from that of bone-fide HSCs from human tissues, we looked at cell-type specific genes curated from the literature (**Figure 3D**). HSC-specific genes, such as GATA2, MIF, DDX21, and CDK6, are expressed solely in both the *bone fide* HSC populations and are gradually upregulated as iPSC differentiation cultures commit to the HSC fate. These genes show low expression or no expression at all in the endothelial or AGM-like clusters. On the contrary, these clusters have high expression of endothelial-specific genes, such as PECAM1, ID3, and Sox17. Moreover, MPP and blood cell populations begin to downregulate genes enriched in *bone fide* and iPSC derived HSCs, while upregulating lineage-specific genes, such as GPX1, LYZ, CD14, and CD3D. To confirm that these gene sets are enriched within the given cell population, we utilized Seurat’s module to score sets of genes (**Figure 3E**). This analysis reveals enrichment of HSCs genes in both the *bone fide* and iPSC derived HSCs with no expression of endothelial genes. Additionally, endothelial gene sets were only enriched in the endothelial and AGM-like clusters. This is further exemplified by overlaying cell-type specific genes onto the UMAP clusters (**Figure 3D**). We find that PECAM1, KDR, CDH5, and SOX17 are strongly enriched in clusters that we have annotated as endothelial or AGM-like hemogenic endothelium. As SOX17 has been shown to mark the emergence of hemogenic endothelial (HE)^23^ and in the process, regulate the HOXA locus^24^, these data provide further evidence that our differentiation protocol faithfully directs iPSCs towards a definitive HSC fate that is intra-embryonic through a HE intermediate. Additionally, CD34 expression was found in all clusters, however, only pre-HSC, HSC, MPP, and blood cells clusters express PTPRC (CD45), most likely owing to the fact that an EHT event has occurred where nascent HSCs begin to bud off the adherent HE. Moreover, HOXA9 expression has been shown to be indicative of a definitive HSC program^11^. We find that HOXA9 is expressed early on in differentiation in the HE and AGM-like clusters, and its expression is sustained in the HSC clusters. In contrast, HSC-specific genes, such as MYB and MYC, are only expressed in the pre-HSC clusters with no detectable expression in HE and AGM-like clusters. Thus far, gene expression patterns suggest that after mesoderm induction of iPSCs, cells undergo stage-specific developmental transitions to a HE population through an AGM-like stage to become definitive HSCs.

To further confirm our hypothesis, we performed RNA velocity analysis using scVelo^25^ of our datasets to elucidate whether our directed differentiation faithfully recapitulates our inferred developmental trajectories (**Figure 3G**). This analysis not only considers gene expression, but it uses the kinetics of splicing patterns to draw developmental trajectories, therefore is a more reliable method for understanding our *in vitro* developmental processes. We find that trajectories drawn begin at the endothelial stage and feed into the AGM-like hemogenic endothelial. Moreover, the trajectories also suggest that the AGM-like hemogenic endothelial cells give rise to both the adherent pre-HSC, HSC, MPP, and blood cell clusters, although not directly to the pre-HSC cluster. However, there is a direct trajectory from the pre-HSC cluster to that of the HSC and its more differentiated progeny. Interestingly, the trajectories drawn from the iPSC derived pre-HSC cluster also include inferred trajectories toward bone-fide HSCs.

Taken together, these data suggest that our directed differentiation protocol faithfully recapitulates the developmental steps *in vitro* that are required to direct iPSCs to a definitive HSC through a definitive AGM-like hemogenic endothelium.

### HSPCs efficiently generate myeloid cells including polarized macrophage

Evidence indicates spatial, phenotypical, and functional differences in the successive contributions of myeloid cells originating from primitive through definitive HM^26^. Human macrophages are extremely plastic *in vitro*, thus capable of polarization to an M1 or M2 phenotype upon cytokine exposure^27^. While these states may not exactly represent *in vivo* counterparts, the model can be extremely useful to examine pro- and anti-inflammatory programs in macrophages for disease modeling. Macrophages were subjected to *in vitro* polarization from an M0 state to either an M1 or M2 state to further tease out differences between primitive and definitive macrophages. We sought to recapitulate these molecular differences between primitive iPSC derived hematopoiesis protocol^28^ with macrophage generated from the AGM-like definitive HPSCs and their ability to make and polarize macrophage *in vitro* in comparison to primary human monocyte derived macrophage (**Figure 4A**).

**Figure 4:**
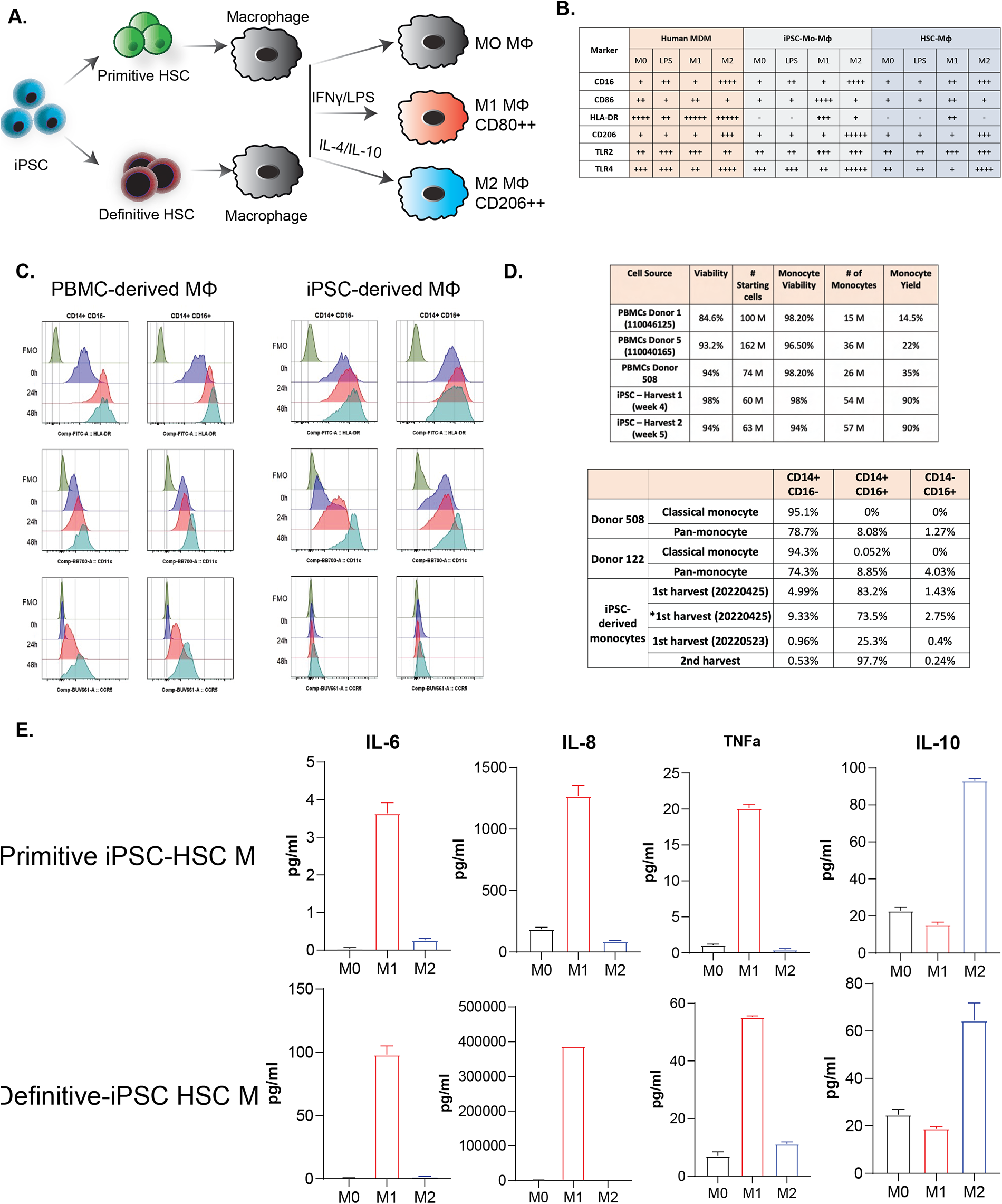
A. Schematic of differentiation approach to testing the differences between our iPSC derived definitive HSCs with that of published primitive protocols with and without polarizing agents. B. Macrophages were differentiated from human monocytes (CD14+), iPSC derived HSCs, and HSCs from CD34+ umbilical cord blood. The table summarizes the expression of M0 (naïve) and polarized macrophage (M1 and M2) markers assessed by flow cytometry. C. Flow cytometry plots comparing the expression of classical and polarized macrophages from PBMCs and iPSCs. D. Table summarizing the yield, donor variability, and percentage of both pan- and classical-monocyte markers. E. MSD cytokine analysis of M0, M1, and M2 polarized macrophages from primitive and definitive iPSC derived HSCs.

All three populations of macrophage were examined by flow cytometry marker expression of CD16, CD86, HLA-DR, CD206, TLR2, and TLR4 and striking similarities were observed between them apart by the lower expression of HLA-DR on the iPSC derived macrophage (**Figure 4B**). PBMC and definitive iPSC derived monocytes had similar flow cytometry profiles when cultured *in vitro* (**Figure 4C**). Cytokine profiling was conducted using meso-scale discovery electrochemiluminescence (MSD) to demonstrate both primitive and definitive iPSC derived macrophage upregulated either the proinflammatory cytokines IL-6, IL-8, and TNFa, or the anti-inflammatory cytokine IL-10 in response to M1 and M2 polarization, respectively (**Figure 4E**).

### HSPCs can maintain gene edits, including the delivery of a functional HER2 chimeric antigen receptor

One strength of human iPSCs is that they are amendable to genome engineering and subclones can be created using lentiviral gene delivery to generate cell lines. Human iPSC were transduced using a commercially available CAR consisting of a Trastuzumab derived scFv-CD28-41BB-CD3ζ targeting the breast cancer antigen erb-b2 receptor tyrosine kinase 2 (ERBB2/HER2)^29^ to evaluate genome silencing during the differentiation to definitive HPSCs, a technical hurdle often encountered during the process of iPSC directed differentiation (**Figure 5A**). To overcome these hurdles, we chose a lentiviral vector that drives the expression of the HER2 CAR a strong promoter that is not prone to silencing, such as EF1A^30^. Additionally, we chose a high multiply infection (MOI) of 25 to ensure enough integrations of the vector into the iPSCs to overcome any heterochromatinization that may occur during differentiation. First, we evaluated the expression of the CAR on the cell surface of iPSCs by incubating the cells with a FITC-conjugated HER2 antigen consisting of amino acids threonine 23-threonine 652. Flow cytometric analysis of HER2 antigen binding to CAR-transduced iPSC revealed that most cells express the CAR on the cell surface (**Figure 5B**). To rule out nonspecific binding of the antigen, untransduced cells were incubated with the HER2 FITC-conjugated antigen. There was no discernable FITC signal detected compared to unstained CAR iPSC lines. To assess whether the expression of the HER2 CAR is sustained after differentiation, we subjected the HER2 CAR iPSC lines toward a definitive HSC using our directed differentiation protocol. We find that robust expression of the HER2 CAR is found on HSPCs derived from our HER2 CAR iPSC lines (**Figure 5C**). However, we find that there is a drop in differentiation efficiency where the percentage of untransduced LT iPSC HSPCs yield 28.7% CD34+CD45+ compared to the 16.0% yield from the HER2 CAR iPSC lines.

**Figure 5:**
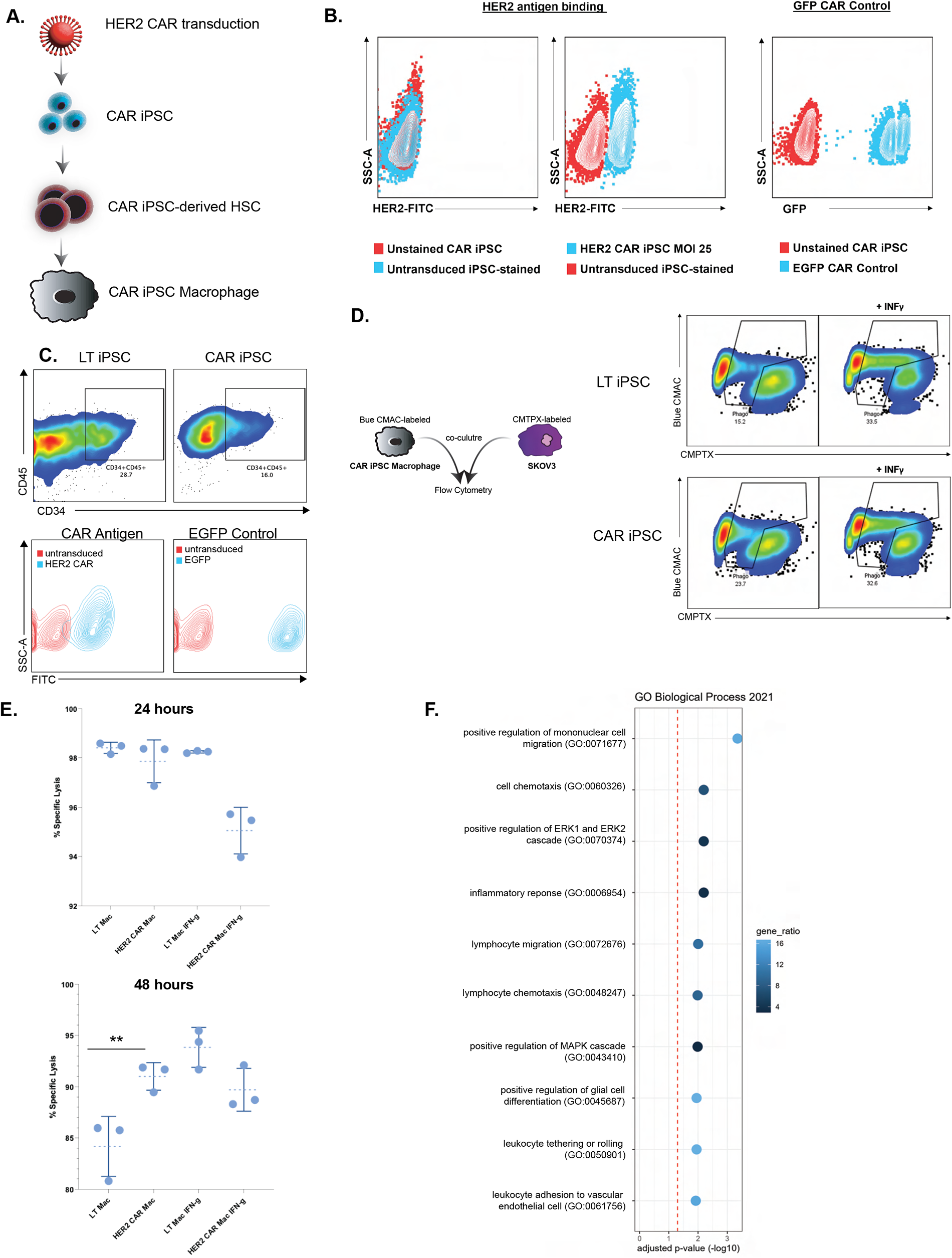
A. Experimental approach for the generation of HER2 CAR macrophages by lentiviral transduction of the CAR in iPSCs B. HER2 antigen binding in transduced HER2 CAR iPSC and the GFP CAR control. Unstained and stained iPSCs that do not have the CAR insertion were used as negative controls for unspecific antigen binding. C. Representative flow plots showing expression of CD34+ and CD45+ after untransduced and HER2 CAR transduced iPSC lines were differentiated to definitive HSC. Expression of the CAR was confirmed by HER2 antigen binding. D. The efficacy of iPSC derived untransduced and HER2 CAR transduced macrophages to phagocytosis SKOV3 HER2+ cancer cells was assessed by labeling macrophages with DAPI (CMAC) and SKOV3 (CMTPX) with TRICTC Cell Tracker dyes. Cells were mixed at a 10:1 (Effector: target) ratio for 24 and 48 hours. Doubles-positive cells were assessed by flow cytometry. E. Quantification of Figure D by calculation percent specific lysis. F. Enrichr gene ontology plot of the biological process of genes that are upregulated in iPSC derived HER2 CAR M1 macrophages cultured with SKOV3 versus HER2 CAR M1 macrophages. Approximately 98 genes were upregulated with a p-value < 0.05 and log2FC > 1.

To test the functionality of these HER2 CAR HSPCs, we directed their differentiation toward macrophages and assessed their ability to phagocytose their target *in vitro*. To do so, we co-cultured SKOV3 cells that have high expression of HER2 with HER2 CAR macrophages. Macrophages were labeled with a blue CMAC cell tracker dye, whereas SKOV3 cancer cells were labeled with CMPTX. Cells were then mixed at an effector to target (E:T) of 10:1, where the effort is the CAR macrophage and the target SKOV3. As the macrophages begin to engulf their target, they become positive for CMPTX as the cancer cell is internalized. Analysis by flow cytometry revealed that iPSC derived macrophages were 23.7% positive for both CMPTX and CMAC blue compared to the 15.2% (**Figure 5D**). Additionally, we used interferon-gamma as a positive control as it has been shown to increase macrophage phagocytosis^31^. To carefully quantify these phagocytic events in a systematic manner, we calculated the percent specific lysis as discussed here^32^, by measuring the CMPTX MFI of the co-cultured samples, SKOV3 alone, and macrophages alone. Interestingly, at 24 hours post-co-culture, there is no significant difference between untransduced macrophages compared to CAR macrophages. However, there is a significant increase in phagocytosis at 48 hours post-co-culture comparable to that of interferon-gamma-treated untransduced LT macrophages, suggesting a time-dependent event that does not require macrophage polarization (**Figure 5E**).

As the CAR we utilized contains a CD3ζ intracellular signaling domain, we sought to understand how this signaling influences the transcriptional response of iPSC derived macrophages upon engaging the antigen *in vitro*. First, we polarized our iPSC derived macrophages towards an M1 phenotype and either cultured them alone or with SKOV3 cells. The cultures were then harvested for RNA extracted and submitted for bulk RNA sequencing. We then ran differential expression analysis on the HER2 CAR M1 macrophages cultured with SKOV3 versus HER2 CAR M1 macrophages alone. We found 98 upregulated genes with a p-value less than 0.05 and a log2 fold-change greater than one. Gene ontology of biological process reveals that genes upregulated in HER2 CAR M1 macrophages with SKOV3 are involved in cell migration, ERK 1/2 signaling, and inflammatory response (**Figure 5G**). Upregulation of genes involved in cell migration in the presence of SKOV3 suggests that the HER2 CAR directs the macrophage to its target for phagocytosis. Furthermore, we find classic hallmarks of CAR activation in our HER2 CAR macrophages exemplified by upregulated genes involved in the positive regulation of ERK1 and ERK2 cascade, which has been shown to be downstream of CD3ζ TCR signaling^33^.

### HSPCs can differentiate into T-cells via an artificial thymic organoid and multilineage blood in vivo

The iPSC derived definitive HPSCs or CD34 positive human cord blood (CB) cells were aggregated with MS5-DLL1 feeder cells in order to compare the T-cell potential between the two populations by an artificial thymic organoid (ATO) method recently described^34^ (**Figure 6A, B**). While the human cord blood cells were three-fold more effective in making CD3/TCRab positive cells the iPSC derived definitive HPSCs could make T-cells (**Figure 6C**), and when gating on CD3, the majority of the cells were CD8 single positive, in contrast to the majority of CB cells producing CD4/8 double positive T-cells (**Figure 6D**).

**Figure 6:**
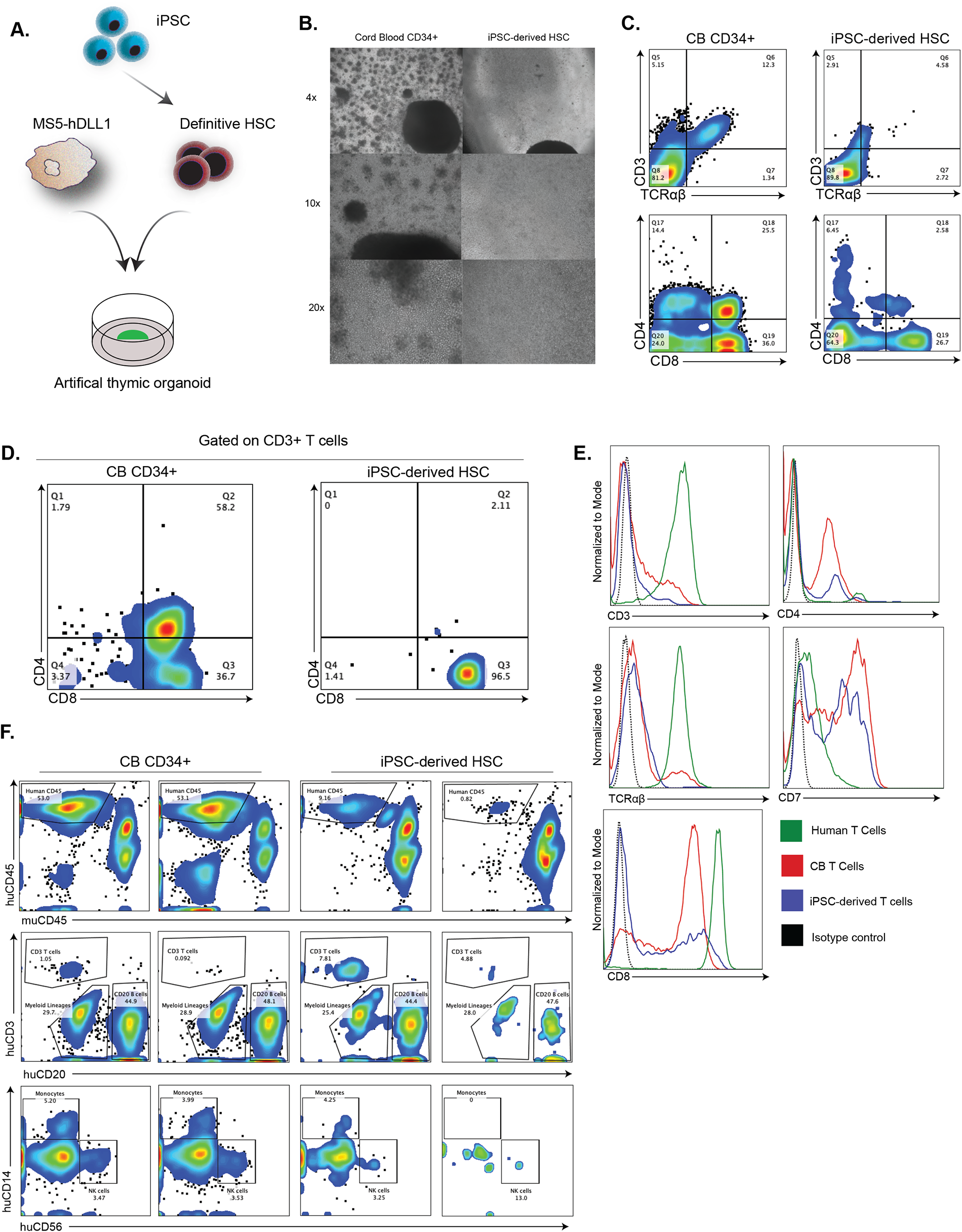
A. Experimental approach for the generation of T-cells from iPSC derived definitive HSCs using artificial thymic organoids (ATO). B. Brightfield images at various magnifications of the same field after 4 weeks of differentiation to T-cells using both cord blood and iPSC derived HSCs in ATOs. C. Representation flow plots showing the expression of CD3 and TCRab double positive populations (top), along with CD4 and CD8 expression (bottom) from ATO differentiations gated on the total number of cells. D. Same plots as Figure 6C, except gates were set to only include cells that were CD3+. Expression of CD4 and CD8 were assessed within the total CD3+ population. E. Flow cytometry flow showing the expression of classical T cell markers, such as CD4, TCRab, CD7, and CD8 gated on total viable cells. Expression patterns of these cell surface markers were compared between cord blood to iPSC derived HSCs in the ability of these cells to give rise to T cells in ATO cultures. F. Flow cytometry of peripheral blood from NSG-SGM3 immunodeficient mice grafted with either CD34+ cord blood (CB) or iPSC derived HSCs at 12 weeks post-injection. Multilineage engraftment was observed (huCD45), including the presence of T-cells (huCD3), B-cells (huCD20), monocytes (huCD14), and NK-cells (huCD56) based on their respective cell surface markers. Two out of five animals grafted with the iPSC derived HSCs are shown and had iPSC derived peripheral blood contribution.

Robust long-term multilineage blood reconstitution in a conditioned animal model is one functional standard for defining an LT-HSC. We sought to examine the homing and engraftment potential of our iPSC derived definitive HPSCs in comparison to CB cells, known to contain LT-HSCs. iPSC derived definitive HPSCs and CB cells were enriched for CD34 and injected into irradiated immunodeficient NOD triple-transgenic mice expressing human IL3, GM-CSF (CSF2) and SCF (KITLG), called NSG-SGM3. The peripheral blood of the NSG-SGM3 mice was examined at 12 weeks post-transplant for both the level of engraftment by human CD45 and multilineage reconstitution using a panel of markers to detect B-cells, T-cells, NK-cells, and monocytes. In four of five CB cell recipients, robust multilineage reconstitution was observed (**Figure 6E**), and one animal lacked robust T-cell contribution. In comparison, two of the five animals transplanted with iPSC derived definitive HPSCs greater than 1% multilineage engraftment was observed (**Figure 6F**).

## Discussion

One of the key strengths of human pluripotent stem cells is having control over the timing and order of human development. This enables researchers to observe and understand aspects of early human development otherwise not observed due to access to primary tissue. Capturing a high efficiency definitive AGM-like hematopoiesis from pluripotent stem cells *in vitro* has been an elusive endeavor. We have been able to observe and demonstrate four discrete phases of development, including mesoderm specification, endothelium specification, EHT, and HPSC expansion and differentiation on the basis of morphology, flow cytometry marker expression, qRT-PCR, single cell sequencing, and functional analysis *in vitro* and *in vivo*.

Definitive hemogenic endothelium arises from an intraembryonic mesoderm lineage, which is marked by the classical T-box transcription factor TBXT. Moreover, in vitro studies of directed differentiation of iPSCs to an intraembryonic mesodermal lineage have been isolated using the cell surface markers CXCR4 and KDR/FLK1^22^. Interestingly, these double-positive cells have been shown to be retinoic acid-dependent in their ability to give rise to definitive HSCs, whereas CXCR4 single-positive cells are retinoic acid-independent and primarily give rise to extraembryonic hematopoietic lineages. However, these studies were done in a 3-dimensional embryoid body (EB) differentiation format that creates signaling gradients. With the use of our TBXT-EGFP reporter, we uncover an additional layer of information as this gene instructs the formation of the primitive streak during gastrulation where intraembryonic mesoderm is specified (REF). Its activity has not been reported to be observed in the formation of extraembryonic mesoderm (REF). We observe the transcriptional activity of TBXT in both the CXCR4+KDR/FLK1- and CXCR4+KDR/FLK1+ populations. Our data suggest that both these populations can give rise to intraembryonic mesoderm in our directed differentiation protocol. Due to the limitations of our experimental design, we believe further investigation is warranted to decipher the developmental origins of these populations. We believe that our differentiation protocol allows us to further investigate these remaining questions due to the 2-dimensional nature allowing us to follow differentiation dynamics more faithfully compared to an EB format.

In contrast to several other reports where hematopoietic progenitor and stem cells have been derived from pluripotent stem cells, we observe the emergence of a RUNX1C+ hematopoiesis up to a week later, between days 14 and 21. The delay in EHT correlates with the later developmental timing of an emergence of a hemogenic endothelium from the AGM and results in a much higher HPSC potency of one in five sorted cells being a CFU as measured by limiting dilution methylcellulose. The higher efficiencies translate into a robust system for generating mature blood cell fates from the iPSC derived HPSCs. One such indicator is the copious production of myeloid and lymphoid cells *in vitro*. Monocytes and macrophage production are particularly productive protocols wherein we demonstrated the capability of gene editing within the iPSC derived hematopoietic system, despite iPSCs being notorious for silencing genetic material and only a limited number of promotors function upon differentiation of the cells^30^.

One hallmark of a definitive hematopoiesis from iPSCs is increased and more balanced differentiation of lymphocytes from HPSCs^14^. The ATO and *in vivo* studies are further evidence our iPSC derived HPSCs are definitive hematopoietic progenitors capable of lymphopoiesis. The *in vivo* differentiation results are particularly compelling in that they produce a comparable repertoire of B-, T-, and NK-cells and persist in the NSG-SGM3 animal model upwards of 12 weeks when compared to human CB cells.

## Materials and Methods

### Directed differentiation of endothelium and HPSCs from iPSCs

For HPSC differentiation, cells were rendered to single cells using Accutase (ICT) and replated on PLO/Laminin treated tissue culture plates in SFD media (IMDM:Ham’s F12, Bovine Albumin, B27, N2, glutamax, Pen/strep, L-ascorbic acid, monotrioglycerol and holo-transferin) containing Y-27632, 10ng/ml BMP4, and 25ng/ml bFGF. On day 1, SFD media was added containing 8uM CHIR99021, 10ng/ml BMP4, and 25ng/ml bFGF. CHIR99021 and BMP4 were withdrawn on day 3 and 25ng/ml of VEGF was added. The cells were fed from days 6-8 with StemPro34 (Thermo) containing 12.5 ng/ml bFGF, 25ng/ml VEGF, 50ng/ml SCF, 25ng/ml IL-6, 25ng/ml IL-3, 25ng/ml FLT3L, 5ng/ml IL-11, and 2U/ml EPO, and from days 9-13 with the prior media containing 10ng/ml BMP4, 10ng/ml SHH, 10ug/ml Angiotensin II, and 100uM Losartan K. Finally, on days 14-17 the cells were fed with StemPro34 containing SCF, 25ng/ml IL-6, 25ng/ml IL-3, 25ng/ml FLT3L, 5ng/ml IL-11, and 2U/ml EPO.

### Flow cytometry and immunofluorescence and colony forming units

HPSCs as defined by positive for CD34, CD45, CD90 and negative for CD38 and a lineage cocktail described above were sorted into a 96-well plate containing human methylcellulose in a limiting dilution of 20, 10, 5, 2, or 1 cell. The 96-well plates were placed in a tissue culture incubator at 37 degrees for 14-21 days and CFC frequency was calculated based on established methods^35^.

### Cell line engineering

The TBXT and RUNX1C EGFP lines were generated by Alstem. The donor vectors consisted of right and left homology arms containing a P2A-H2B-EGFP followed by an excisable puromycin selection cassette. After transfection, cells were then selected with 0.25μg/mL puromycin for 5 days. Subsequently, cells were plated as single cells for clone selection. PCR amplification was used to detect correctly targeted positive clones for expansion. Screening primers can be found in table 1.

HER2 CAR lentiviral particles were purchased from CreativeBio Labs (CAR-C1) consisting of scFv-CD28-41BB-CD3ζ. HER2 CAR LT iPSC lines were generated by lentiviral transduction. Briefly, LT iPSC cultures were treated with Accutase to obtain a single-cell suspension. Approximately 2×10^5^ cells were plated on Matrigel-coated plates in StemFlex media containing 1μg/mL polybrene. Lentiviral particles were then added to the cell suspension at a multiplicity of infection of (MOI) 25. Cultures were then incubated overnight. To select for cells that received the construct, 1μg/mL puromycin was added to the media for 48 hours. Afterward, surviving clones were treated with Accutase to obtain a single-cell suspension and replated in media containing puromycin for an additional 48 hours. To assess the expression of the HER2, approximately 1×10^5^ cells were incubated with 2.5 μg of FITC conjugated HER2 (Acro Biosystems HE2-HF256) and analyzed by flow cytometry.

### HPSC single-cell sequencing and analysis

Single-cell RNA-seq was performed using the either the Chromium 3’ V2 or Chromium 3’ V3 chemistry (10X Genomics). Live cells for each sample were sorted and added to reverse transcription master mix with the appropriate volume of RNAse-free water per manufacturer’s guidelines and encapsulated with gel beads using partitioning oil to generate Gel Bead-in-EMulsions (GEMs). After encapsulation, barcoded mRNAs were extracted from GEMs and cleaned. 11 cycles of cDNA amplification were performed using standard temperature settings according to manufacturer’s recommendations. Completed libraries were pooled and sequenced on Illumina NovaSeq6000 S4 flow cells using the sequencing cycle specifications of 26:8:0:98 (Read1:i7 index:i5 index:Read2) for Chromium 3’ V2 or 28:8:0:91 for Chromium 3’ V3. Sample-specific FASTQ files were generated from raw Illumina BCL files using bcl2fastq.

FASTQ reads were aligned against human reference genome GRCh38-3.0.0 (Cellranger software v3.0.2, 10X Genomics). Raw count matrices of barcoded, mapped reads were then further filtered using a multi-step process to get rid of empty droplets and outlier transcriptomes. High-quality transcriptomes were then defined as cell barcodes having >1000 features detected, <10% fraction of mitochondrial RNA reads and a sequence saturation > 40%. The predicted cell cycle phase for each cell was also determined using the machine learning-based approach described by Scialdone et al^36^. All libraries were normalized using the procedure outlined in Ellwanger, et al^37^. The cell-based size factors were normalized between libraries based on the ratio of average UMI counts between libraries using the R package batchelor. The resulting UMI count matrix was log2-transformed and used for downstream analyses.

To quantitatively find the gene expression features that have significantly high variance between cells, we used the approach implemented in Ellwanger, et al^37^. Only genes with expression in >3% of cells were considered. The variance of each gene was fitted as a function of mean expression. Genes were tested for significance using the R package scran by modeling the residuals with an F-distribution, and genes with a p-value < 0.001 and a minimum average expression ≥1 were considered significant. To ensure that informative variable features were not being excluded, variable feature identification, dimension reduction, and clustering were performed using genes with minimum average expression >0.1 and no novel cell clusters were identified.

Data dimensionality was reduced using Diffusion Maps (Coifman & Lafon, 2006), since this algorithm can resolve both linear and non-linear substructures across cell states in a deterministic manner. Using the plotExplanatoryVariables function in the R package scater (version 1.0.4), library batch and chemistry-specific (i.e. 10X 3’ V2 versus 10X 3’ V3) variations in cellular expression were detected and determined to be the major confounding variables masking biological variation. Therefore, these factors were regressed out during dimension reduction to increase the signal-to-noise ratio of the spectral embedding. After factor analysis using a scree plot to determine the optimal number of diffusion components to describe variance within the dataset, we used the first 45 components to compute a two-dimensional uniform manifold approximation and projection of single cells.

To assign clusters (cell states), a shared Nearest Neighbor (SNN) graph was first constructed of the top 45 diffusion components using the R package Seurat^38^. Subsequently, the Louvain algorithm implemented in Seurat was used to cluster cells.

To understand the differentiation dynamics of the dataset, RNA velocity was calculated using scvelo package in python with dynamic modeling and a minimum mass of 3.

### Macrophage differentiation, polarization, MSD, and phagocytosis assay

The monocytic lineage differentiation protocol followed a previously established hematopoietic differentiation protocol^28^. The protocol consists of 5 sequential steps by which mature macrophages are differentiated from CD14+ monocytes. In brief, monocytes were cultured in X-vivo 15 medium with X-VIVO 15 with 2 mM glutamax, 55 uM b-mercaptoethanol, 1x B27, 1x N2 and 100ng/ml hMCSF for 7 days, change medium at day 3. For definitive iPSC derived macrophage from HPSCs the same culture medium as monocytes protocol described as above, change medium every 3-4 days and culture for 2 weeks. Macrophages were polarized to M1 phenotype by the culture medium with 25ng/ml IFNg and 10ng/ml LPS for 48hrs. Polarized to M2 phenotype by culture medium with 40ng/ml IL4. All MSD experiments were performed using cell supernatant one the Meso Sector S 600MM according to the manufacturer’s protocols.

The phagocytosis assay is based on a prior study^32^, and the following equation was used: % specific lysis = [(Sample signal − Tumor alone signal)/(Background signal − Tumor alone signal)] × 100.

### T-cell differentiation

The T-cell differentiation follows a recently published study from Montel-Hagen, et al.^34^. In brief, 5×10^5^ MS5-hDLL1 cells were harvested using trypsin and combined with 5×10^3^ of either the iPSC derived definitive HPSCs or CD34 positive human CB cells and in base hematopoietic medium containing EGM2, 10uM Y-27632, and 10uM SB-431542 and centrifuged at 300g. The cell pellet was resuspended in 6 ul fresh hematopoietic induction medium and plated on a 0.4um Millicell trans-well insert. The insert was placed in a well of a 6-well plate containing 1ml of hematopoietic induction medium and media was changed every 2-3 days for 1 week then replaced with media containing hematopoietic cytokines rhTPO 5 ng/ml, rhFLT3L 5 ng/ml, and rhSCF 50 ng/ml. At week 2, T-cell differentiation was induced by changing to RB27 media (RPMI 1640, 4% B27 supplement, 30 µM L-ascorbic acid, 1% Glutamax, 10 ng/ml rhSCF, 5 ng/ml rhFLT3L and 5 ng/ml rhIL-7.

### Animals, LT-HSC engraftment, and analysis

Deidentified umbilical cord blood (UCB) CD34+ cells were purchased from Lonza. iPSC derived HSCs were obtained from cultures at day 22 of differentiation and sorted based on CD34+ by Miltenyi magnetic activated cell sorting (MACS). Four-to five-week-old NSG-SGM3 (NSGS) mice were irradiated at a single dose of 1 Gy then conditioned for 4-6 hours. Mice were anesthetized with isoflurane and approximately 3×10^4^ UCB CD34+ and 1×10^6^ iPSC derived cells in 100μL of PBS were delivered via the retroorbital sinus using a 27g insulin needle. Peripheral blood was obtained by terminal cardiac puncture. Blood was then lysed using ACK lysis buffer prior to staining for flow cytometric analysis. Staining cocktail consisted of BV421 CD3, PE CD11c, BUV496 CD14, FITC CD20, APC-R700 CD33, BUV805-CD56, PE-Cy7 human CD45, and APC murine CD45. Cell viability was determined by 7-AAD.

## Acknowledgements

We’d like to thank Menno Van Lookeren Campagne, Zhifei Shao, John Ferbas, and Amgen Cytometry Sciences for their technical knowledge and scientific support.

---

**Figure S1.**
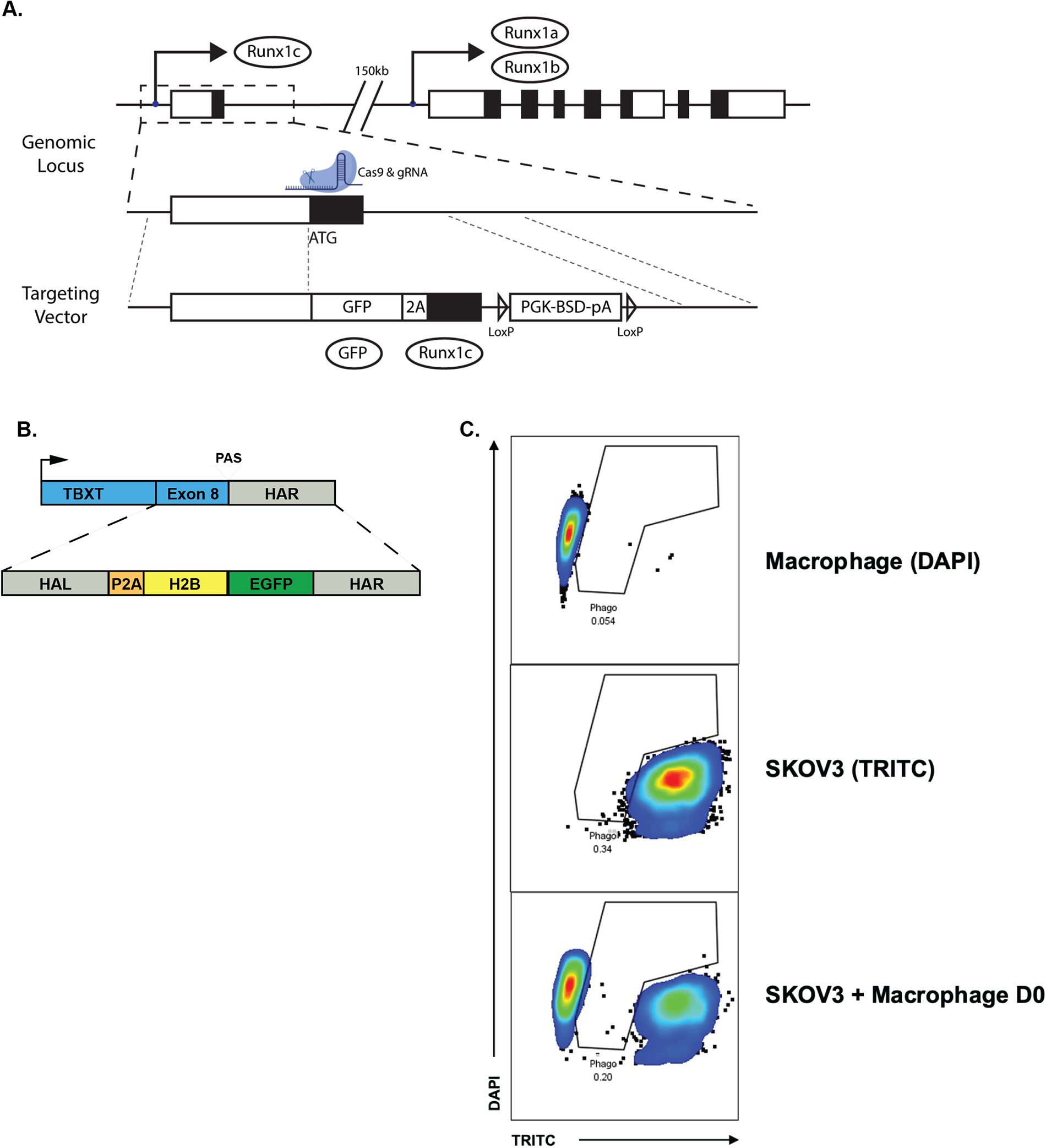
A. Targeting strategy for generating RUNX1c reporter iPSC lines. B. Targeting strategy for the generation of TBXT EGFP LT iPSC reporter line C. Control flow plots showing DAPI and TRITC fluorescence with the specified cell types without co-culture. Day 0 (D0) cultures were trypsinized and mixed immediately prior to acquisition.

**Figure S1.**
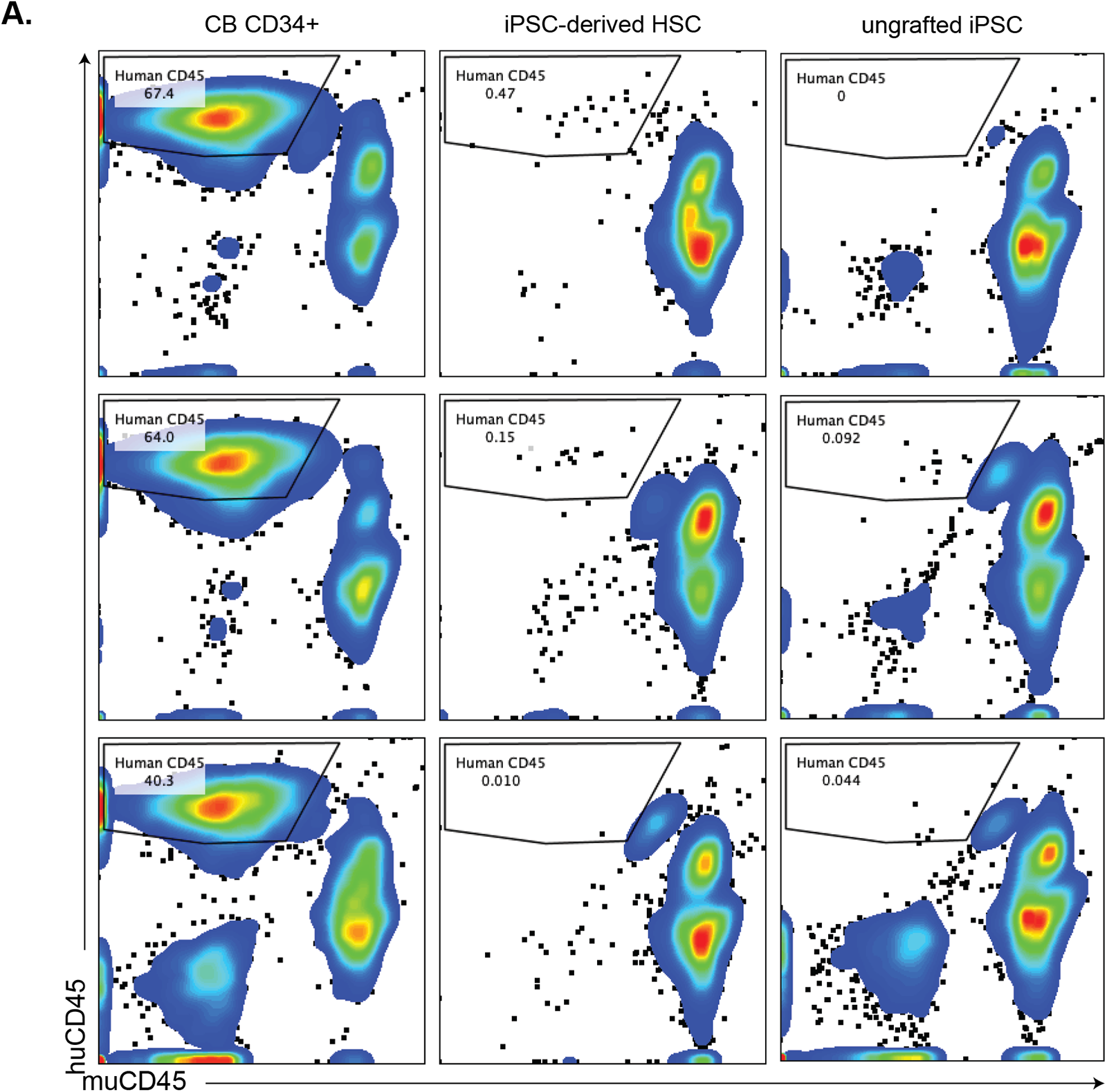
A. Flow cytometry plots of peripheral blood from remaining NSG-SGM3 within the experimental cohort. Chimerism was assessed using a human specific CD45 antibody (clone H130). Animal cohorts include those grafted with 3x10^4^ umbilical cord blood CD34+ cells (left), freshly cultured iPSC-derived HSCs (middle), and lastly iPSC-derived HSCs that were cryopreserved prior to engraftment (right).

**Supplemental Table 1.**
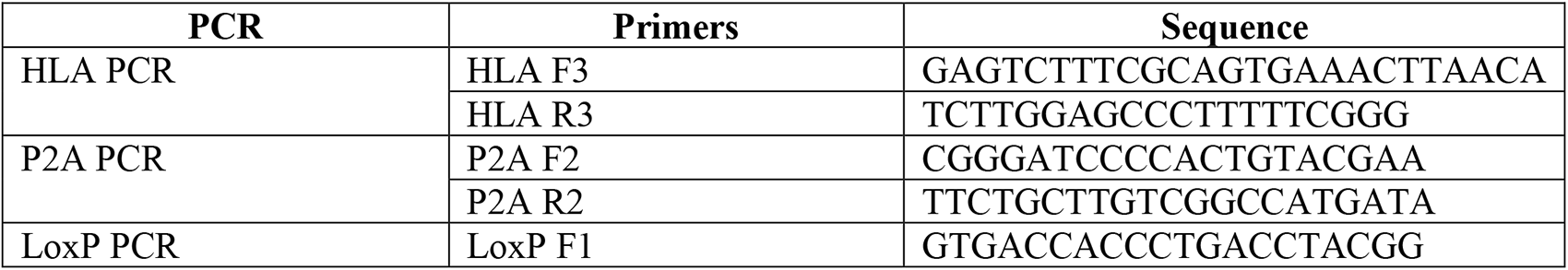
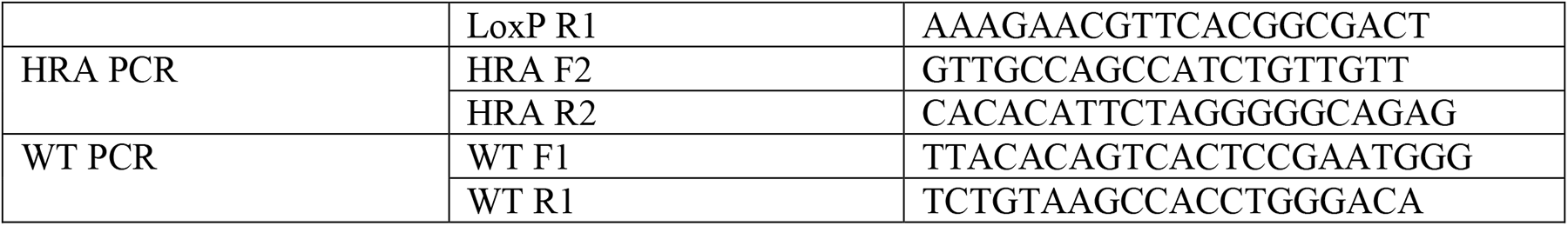

## References

1. Takahashi, K. et al. Induction of pluripotent stem cells from adult human fibroblasts by defined factors. Cell 131, 861–872 (2007).

2. Thomson, J.A. et al. Embryonic stem cell lines derived from human blastocysts. Science 282, 1145–1147 (1998).

3. Notta, F. et al. Isolation of single human hematopoietic stem cells capable of long-term multilineage engraftment. Science 333, 218–221 (2011).

4. Canu, G. & Ruhrberg, C. First blood: the endothelial origins of hematopoietic progenitors. Angiogenesis 24, 199–211 (2021).

5. Daniel, M.G., Pereira, C.F., Lemischka, I.R. & Moore, K.A. Making a Hematopoietic Stem Cell. Trends Cell Biol 26, 202–214 (2016).

6. Suzuki, N. et al. Generation of engraftable hematopoietic stem cells from induced pluripotent stem cells by way of teratoma formation. Mol Ther 21, 1424–1431 (2013).

7. Elcheva, I. et al. Direct induction of haematoendothelial programs in human pluripotent stem cells by transcriptional regulators. Nat Commun 5, 4372 (2014).

8. Duan, F. et al. Biphasic modulation of insulin signaling enables highly efficient hematopoietic differentiation from human pluripotent stem cells. Stem Cell Res Ther 9, 205 (2018).

9. Sugimura, R. et al. Haematopoietic stem and progenitor cells from human pluripotent stem cells. Nature 545, 432–438 (2017).

10. Zhu, Y. et al. Characterization and generation of human definitive multipotent hematopoietic stem/progenitor cells. Cell Discov 6, 89 (2020).

11. Calvanese, V. et al. Mapping human haematopoietic stem cells from haemogenic endothelium to birth. Nature 604, 534–540 (2022).

12. Sturgeon, C.M., Ditadi, A., Awong, G., Kennedy, M. & Keller, G. Wnt signaling controls the specification of definitive and primitive hematopoiesis from human pluripotent stem cells. Nat Biotechnol 32, 554–561 (2014).

13. Ng, E.S. et al. Differentiation of human embryonic stem cells to HOXA(+) hemogenic vasculature that resembles the aorta-gonad-mesonephros. Nat Biotechnol 34, 1168–1179 (2016).

14. Kennedy, M. et al. T lymphocyte potential marks the emergence of definitive hematopoietic progenitors in human pluripotent stem cell differentiation cultures. Cell Rep 2, 1722–1735 (2012).

15. Okuda, T., van Deursen, J., Hiebert, S.W., Grosveld, G. & Downing, J.R. AML1, the target of multiple chromosomal translocations in human leukemia, is essential for normal fetal liver hematopoiesis. Cell 84, 321–330 (1996).

16. Challen, G.A. & Goodell, M.A. Runx1 isoforms show differential expression patterns during hematopoietic development but have similar functional effects in adult hematopoietic stem cells. Exp Hematol 38, 403–416 (2010).

17. Navarro-Montero, O. et al. RUNX1c Regulates Hematopoietic Differentiation of Human Pluripotent Stem Cells Possibly in Cooperation with Proinflammatory Signaling. Stem Cells 35, 2253–2266 (2017).

18. Chambers, S.M. et al. Highly efficient neural conversion of human ES and iPS cells by dual inhibition of SMAD signaling. Nat Biotechnol 27, 275–280 (2009).

19. Chambers, S.M. & Studer, L. Cell fate plug and play: direct reprogramming and induced pluripotency. Cell 145, 827–830 (2011).

20. Bennett, D. et al. Observations on a set of radiation-induced dominant T-like mutations in the mouse. Genet Res 26, 95–108 (1975).

21. Showell, C., Binder, O. & Conlon, F.L. T-box genes in early embryogenesis. Dev Dyn 229, 201–218 (2004).

22. Luff, S.A. et al. Identification of a retinoic acid-dependent haemogenic endothelial progenitor from human pluripotent stem cells. Nat Cell Biol 24, 616–624 (2022).

23. Nobuhisa, I. et al. Sox17-mediated maintenance of fetal intra-aortic hematopoietic cell clusters. Mol Cell Biol 34, 1976–1990 (2014).

24. Jung, H.S. et al. SOX17 integrates HOXA and arterial programs in hemogenic endothelium to drive definitive lympho-myeloid hematopoiesis. Cell Rep 34, 108758 (2021).

25. Bergen, V., Lange, M., Peidli, S., Wolf, F.A. & Theis, F.J. Generalizing RNA velocity to transient cell states through dynamical modeling. Nat Biotechnol 38, 1408–1414 (2020).

26. McGrath, K.E., Frame, J.M. & Palis, J. Early hematopoiesis and macrophage development. Semin Immunol 27, 379–387 (2015).

27. Vogel, D.Y. et al. Human macrophage polarization in vitro: maturation and activation methods compared. Immunobiology 219, 695–703 (2014).

28. Yanagimachi, M.D. et al. Robust and highly-efficient differentiation of functional monocytic cells from human pluripotent stem cells under serum- and feeder cell-free conditions. PLoS One 8, e59243 (2013).

29. Swain, S.M., Shastry, M. & Hamilton, E. Targeting HER2-positive breast cancer: advances and future directions. Nat Rev Drug Discov 22, 101–126 (2023).

30. Norrman, K. et al. Quantitative comparison of constitutive promoters in human ES cells. PLoS One 5, e12413 (2010).

31. Jorgovanovic, D., Song, M., Wang, L. & Zhang, Y. Roles of IFN-gamma in tumor progression and regression: a review. Biomark Res 8, 49 (2020).

32. Klichinsky, M. et al. Human chimeric antigen receptor macrophages for cancer immunotherapy. Nat Biotechnol 38, 947–953 (2020).

33. Hwang, J.R., Byeon, Y., Kim, D. & Park, S.G. Recent insights of T cell receptor-mediated signaling pathways for T cell activation and development. Exp Mol Med 52, 750–761 (2020).

34. Montel-Hagen, A. et al. Organoid-Induced Differentiation of Conventional T Cells from Human Pluripotent Stem Cells. Cell Stem Cell 24, 376–389 e378 (2019).

35. Ploemacher, R.E., van der Sluijs, J.P., Voerman, J.S. & Brons, N.H. An in vitro limiting-dilution assay of long-term repopulating hematopoietic stem cells in the mouse. Blood 74, 2755–2763 (1989).

36. Scialdone, A. et al. Computational assignment of cell-cycle stage from single-cell transcriptome data. Methods 85, 54–61 (2015).

37. Ellwanger, D.C. et al. Prior activation state shapes the microglia response to antihuman TREM2 in a mouse model of Alzheimer’s disease. Proc Natl Acad Sci U S A 118 (2021).

38. Butler, A., Hoffman, P., Smibert, P., Papalexi, E. & Satija, R. Integrating single-cell transcriptomic data across different conditions, technologies, and species. Nat Biotechnol 36, 411–420 (2018).

